# Drosha Regulates Oogenesis and microRNAs Germline Autonomously and Non-autonomously in *C. elegans*

**DOI:** 10.1101/2020.02.26.966648

**Authors:** Amanda L. Minogue, Kenneth A. Trimmer, Jacob H. Seemann, Awdhesh Kalia, Swathi Arur

## Abstract

Small non-coding RNAs regulate multiple aspects of development including germ cell development. The microRNA pathway genes Dicer, Drosha and Pasha have been shown to regulate oocyte meiotic maturation in *C. elegans*. However, Dicer controls oocyte meiotic maturation through endo-siRNAs, rather than microRNAs. A repertoire of Drosha-dependent oocyte-expressed microRNAs were identified which regulate various aspects of oogenesis but not oocyte meiotic maturation. These data lead to the following models: (a) microRNAs function redundantly to regulate oocyte meiotic maturation, (b) Drosha and microRNAs function germline non-autonomously to regulate meiotic maturation. We investigated these models and observed that Drosha regulates oocyte meiotic maturation in a germline non-autonomous manner. Additionally, we uncovered a role for Drosha in regulating pachytene progression and oocyte development in a germline autonomous manner through miR-35 family and miR-51 respectively. Interestingly we also find that though Drosha-dependent oocyte-expressed microRNAs, miR-61 and miR-72, are sufficient to regulate pachytene progression and oocyte development respectively, they are generated in a germline non-autonomous manner. Collectively these data reveal a Drosha-dependent microRNA circuit, which coordinates oocyte development germline autonomously as well as through soma-germline communication.

## INTRODUCTION

Organismal development is tightly controlled by inter and intracellular communication through regulation of molecular and signaling pathways (Perrimon et al., 2012). Small non-coding RNAs such as endogenous small interfering RNA (endo-siRNAs (Storz, 2002)), microRNAs (Hammond, 2015) and piRNAs (Weick and Miska, 2014), among others, are a class of key developmental regulators that communicate intracellularly and traverse tissues to execute their function (Sarkies and Miska, 2014). These small non-coding RNAs range in size from 21 to 30 nucleotides and regulate gene expression by binding to the target mRNA and causing either post-transcriptional degradation or translational inhibition (Storz, 2002); in each case affecting developmental events. Due the small size of these RNAs, they easily traverse through cells and mediate inter-tissue communication (Sarkies and Miska, 2014), and can regulate cellular events in contexts outside of where they are generated. Because of their critical roles in regulating development and the aberrations in their production leading to developmental diseases or cancers (Ardekani and Naeini, 2010; Li et al., 2009; Srinivasan et al., 2013). Therefore, understanding the exact site of generation of these small RNAs and their cell autonomous and non-autonomous roles is critical.

Due to the vast expansion of the regulatory small RNAs in every species it is challenging to attribute function and site of origin *vs* action to each regulator independently (Seitz, 2017). *C. elegans* for example has ∼300 microRNAs, which often function redundantly to execute developmental events (Miska et al., 2007). Thus, to determine the function of these regulators in a given cell or tissue, analysis of pathways that generate small RNAs is conducted by investigating their effects on development. One developmental process which may be uniquely sensitive to small RNA perturbation is oocyte development; which from worms to humans, is largely transcriptionally silent (Greenstein, 2005). As a result, a role for small non-coding RNAs is of great interest in this process; particularly since Dicer was discovered as a regulator of oocyte meiotic maturation in worms (Knight and Bass, 2001). Loss of Dicer, Drosha and Pasha in *C. elegans* each result in the same phenotype: endo-mitotic oocytes due to defects in oocyte meiotic maturation (Knight and Bass, 2001; Rios et al., 2017), resulting in sterile animals. Loss of Dicer from mouse oocytes was also shown to regulate oocyte maturation (Murchison et al., 2007), leading to the model that microRNAs regulate oocyte development and are conserved in their function. However, using tissue specific mosaic analysis, Drake et al. showed that Dicer regulates oocyte meiotic maturation in worms in a germline non-autonomous manner (Drake et al., 2014). Using small RNA sequencing they further went on to show that Dicer regulates oocyte meiotic maturation through the regulation of endo-siRNAs rather than microRNAs. Presumably, endo-siRNAs are generated in the soma and traverse to the germline to regulate meiotic maturation. Following this work, Stein et al. showed that much as in worms, Dicer regulates endo-siRNAs in mice to regulate oocyte maturation (Stein et al., 2015). Further work from Suh et al. in mice suggested that microRNAs are not generated during oogenesis and do not regulate oocyte development (Suh et al., 2010). However, microRNA profiling of dissected germlines from *C. elegans* that contain all of the germ cells and the somatic gonadal cells by Minogue et al. (Minogue et al., 2018) and Bezler et al. led to identification of a very small repertoire of microRNAs (∼7%) in oogenic germlines (Bezler et al., 2019), while Diag et al. (Diag et al., 2018) detected all the microRNAs in the germline. While the difference in how many microRNAs were detected between the two studies could be at the level of sensitivity and specificity in the experiments conducted, nevertheless, these studies suggested that microRNAs are expressed during oogenesis. To determine if these microRNAs are functional, Minogue et al. mapped the localization of mature microRNAs using *in situ* hybridization and functionally tested them with mutant and RNAi based analyses to show that the microRNAs were expressed in developing and arrested oocytes (Minogue et al., 2018). Their analysis showed that loss of microRNAs led to oocyte developmental phenotypes in the region of their expression (Minogue et al., 2018). Surprisingly, none of the microRNA depletions led to oocyte meiotic maturation phenotypes leading to the model that the microRNAs may be redundant in their function to regulate oocyte meiotic maturation, or that germline expressed microRNAs do not regulate meiotic maturation. It also opens up questions as to the exact role of Drosha and Pasha, the microprocessor complex, in oocyte development given that Dicer does not function autonomously in the germline to regulate oogenesis.

To investigate the role of the microprocessor complex during oogenesis and meiotic maturation, we investigated the function of Drosha (which is a critical component of the microprocessor complex) using two deletion alleles that removes the RNAse IIIb domains and germline-specific depletion of Drosha during oogenesis. We observed that Drosha regulates oocyte meiotic maturation in a germline non-autonomous manner. We also investigated whether the generation of Drosha-dependent oocyte-expressed microRNAs is germline autonomous or non-autonomous to mediate oogenesis. Loss of Drosha in the germline specifically results in delays in pachytene progression and oocyte development; these phenotypes are mimicked by loss of the miR-35 and miR-61 microRNAs, which are expressed in late pachytene and into developing oocytes. Surprisingly, however, miR-61 is generated germline non-autonomously for this function; miR-35’s germline autonomous function is sufficient to regulate pachytene progression. Deep RNA Sequencing of miR-35 mutants for potential post-transcriptional targets did not reveal any obvious targets which may regulate chromosomal transition in late pachytene, suggesting that miR-35 may either act through multiple targets or through translational inhibition to regulate this function. In addition, miR-51 and miR-72, which are both expressed only in developing and arrested oocytes, are generated germline autonomously and non-autonomously, respectively, suggesting that similar expression patterns between microRNAs are not sufficient to infer their site of generation. Collectively, our analysis reveals that in the germline Drosha autonomously regulates the oogenic phenotype of chromosomal transition in pachytene through the regulation of the core germline microRNA family, miR-35, and oocyte development through miR-72, while miR-61 and miR-51 are regulated germline non-autonomously for these phenotypes. Oocyte meiotic maturation is regulated in a germline non-autonomous manner through somatic regulation of microRNAs. Together, a Drosha-dependent microRNA circuit coordinates oocyte development autonomously as well as through soma-germline communication.

## MATERIALS AND METHODS

### Experimental model and subject details

Standard *C. elegans* culture conditions at 20 °C were used. The following alleles were used in this study. Linkage group I: *drsh-1(ok369)/hT2G, drsh-1(tm654)/hT2G*. Transgenic lines *drsh-1(ok369)/hT2G* I; bcIs39[(lim-7)ced-1p::GFP+lin-15(+)] V, *drsh-1(tm654)/hT2G* I; bcIs39[(lim-7)ced-1p::GFP+lin-15(+)] V, *drsh-1(ok369)/hT2G* I; ltIs38[pAA1; pie-1::GFP::PH(PLC1delta1) + unc-119(+)] III; ltIs37 [P-pie-1::mCherry::his-58 (pAA64) + unc-119(+)] IV, *drsh-1(tm654)/hT2G* I; ltIs38[pAA1; pie-1::GFP::PH(PLC1delta1) + unc-119(+)] III; ltIs37 [P-pie-1::mCherry::his-58 (pAA64) + unc-119(+)] IV, mkcSi13[sun-1p::rde-1::sun-1 3’ UTR + unc-119(+)] II; *rde-1(mkc36)* V.

### Gonad dissection and analysis

Dissections of adult worms at indicated stage of development to obtain intact gonads were performed (Arur et al., 2011; Arur et al., 2009), under 5 min (immediately after adding levamisole). The dissected gonads were then fixed in 3% paraformaldehyde for ten min, followed by a post-fix in 100% methanol at −20 °C overnight. The fixed and permeabilized gonads were then washed three times with 1X PBS containing 0.1% Tween-20 and processed for immunofluorescence staining as described (Arur et al., 2011; Arur et al., 2009; Gervaise and Arur, 2016; Lee et al., 2007). For the progenitor zone analysis, EdU processing (see below method) was performed before blocking. Germlines were then blocked in 30% NGS for 4 hours. The primary antibody was incubated at room temperature for 4 hours, and secondary antibody was incubated at 4 °C overnight. For the pachytene and oocyte analysis, an average of 10 germlines per genotype per replicate were analyzed with three total replicates performed. For the progenitor zone analysis, 30 germlines were analyzed for each genotype.

### Antibodies

The following antibodies were used for immunofluorescence analysis: rabbit anti-HIM-3 (5347.00.02, Sdix) used at 1:800; mouse anti-phospho-Histone H3 (Ser10) (05-806, Millipore) used at 1:500. Secondary antibodies used were goat anti-rabbit Alexa Fluor 488 (A11008, Life Technologies) used at 1:800; donkey anti-mouse Alexa Fluor 594 (A21203, Life Technologies) used at 1:800. To visualize DNA, the fixed and stained samples were incubated with 100 ng/mL 4′,6′-diamidino-2-phenylindole hydrochloride (DAPI) in 1X PBS with 0.1% Tween-20 for 5 min at room temperature.

### Cell counting

Pachytene cell rows and oocytes were counted manually from single plane images of the germline based on DAPI staining. Apoptotic cells were counted using ImageJ and a custom macro written by Dr. Kenneth Trimmer from Z-stack images. This macro assists in counting cell nuclei from a Z-stack when nuclei may be present in more than one slice.

### EdU labeling

Soaking EdU was performed by washing worms from NGM and OP50 plates three times with M9T (M9 buffer, 0.1% Tween 20) and then the worms were transferred to a flat-bottomed 48-well plate. An incubation with 500 mM EdU solution was performed for 15 min at room temperature in the dark. The animals were then dissected and germlines processed using the Click-iT Plus EdU Alexa Fluor 647 Imaging Kit (ThermoFisher Scientific) per the manufacturer’s recommendations, with minor modifications. Instead of the copper protectant provided with the kit (Component E), 2mM CuSO_4_ (final concentration) was used. Also, the 647 picolyl azide was used at twice the concentration.

### Measuring M phase and S phase indices

The progenitor zone length was measured by counting the number of rows from the distal tip until the onset of either the first HIM-3 positive nucleus or the first crescent shape nucleus. Each nucleus was visualized by DAPI staining. M phase index was calculated as a percent corresponding to the number of pH3 positive cells in the progenitor zone divided by the total number of cells in the progenitor zone. S phase index was calculated as a percent corresponding to the number of EdU positive cells in the progenitor zone divided by the total number of cells in the progenitor zone. The criterion used for distinguishing the progenitors from meiotic cells was the absence of nuclear enrichment of HIM-3 staining. Counting was done using ImageJ and a custom macro written by KAT from Z-stack images.

### Time-lapse imaging

Live animals were placed on a slide and anaesthetized with levamisole. Animals were imaged using a 63X objective every four minutes for 60 to 90 minutes. Z-stacks were acquired utilizing 0.6 μM thick optical sections. Images were taken on a Zeiss Axio Imager upright microscope by using AxioVs40 V4.8.2.0 SP1 micro-imaging software and a Zeiss Axio MRm camera. Approximately 10 animals were imaged per genotype.

### Mating assay

10 wild type males were placed with 3 memb::GFP; mCherry::H2B, *drsh-1(ok369)*; memb::GFP; mCherry::H2B or *drsh-1(tm654)*; memb::GFP; mCherry::H2B L4 hermaphrodites. The mated hermaphrodites were imaged on the Zeiss Axio Imager described above and analyzed for oocyte-to-embryo defects, and the total number of progeny was counted.

### RNA interference (RNAi) by feeding

RNAi was performed by feeding as described previously (Arur et al., 2009). *glp-1* and *drsh-1* RNAi clones (Ahringer RNAi library and Vidal ORFeome Library respectively) were sequenced verified and grown overnight on solid LB agar plates containing 100 μg / ml of Ampicillin and 50 μg / ml of Tetracyclin at 37 °C. Single colonies were then inoculated into LB liquid cultures containing 100 μg / ml of Ampicillin and 50 μg / ml of Tetracyclin at 37 °C and grown to necessary densities as described previously (Arur et al., 2009). The cultures were then seeded onto the standard NGM agar plates supplemented with 1 mM isopropyl β-D-1-thiogalactopyranoside (IPTG) and containing 100 μg / ml of Ampicillin and 50 μg / ml of Tetracyclin. Fresh plates were incubated at room temperature for at least 3 days to allow for bacterial lawn growth. For F2 RNAi, L4 stage animals of the genotype listed were transferred to RNAi plates and allowed to lay progeny for 24 hours. Approximately 3 days later F1 L4s were then transferred to new plates and allowed to lay progeny for 24 hours. Approximately 3 days later F2 L4s were then transferred to new plates and allowed to grow. The F2s were then dissected for analysis at the L4+24h developmental time point. Three replicates were completed for each treatment.

### Quantitative real-time PCR (qRT-PCR)

RNA was collected from indicated strains as well as from RNAi treatments using TRIZOL Reagent (Invitrogen). Three biological replicates, with four technical replicates each, were performed for each genotype for each experiment. Total RNA was purified using the miRNAeasy Mini Kit (Qiagen). Complementary DNA was generated using the iScript cDNA synthesis kit (Bio-Rad Laboratories) from 10 ng total RNA. qRT-PCR analysis was performed using custom primers specific to Actin or *drsh-1*. Expression levels were standardized to Actin and then to Luciferase using the ΔΔC_t_ method of standardization.

### TaqMan assay for detection of mature microRNAs

100 animals for each indicated genotype was collected into TRIZOL Reagent (Invitrogen). Three biological replicates, with four technical replicates each, were performed for each genotype. Total RNA was purified from TRIZOL extraction using the miRNAeasy Mini Kit (Qiagen). Complementary DNA was generated using custom TaqMan assay primers for cel-miR-35-3p, cel-miR-51-5p, cel-miR-61-3p, cel-miR-72-3p, cel-miR-229-5p, and U6 snRNA (Applied Biosystems) from 10 ng of total RNA. qRT-PCR analysis was performed using specific TaqMan assays for each RNA using manufacturers protocol (Applied Biosystems). Expression levels were standardized to the U6 snRNA positive control and then to wild type using the ΔΔC_t_ method for standardization.

### mRNA differential expression analysis

Quality assessment of the NextSeq (Illumina, Inc., San Diego, CA) raw reads sequenced at 40 million read depth was performed with FastQC (Brown et al., 2017; Ward et al., 2019). In preparation for the analysis, low quality reads (average Q-score below 30, more than 4 Ns within read sequence), bases (q score < 10 at the end of the reads), homopolymers (>95% frequency of a particular base), and adapters sequences were removed using a combination of Trimmomatic (Mortazavi et al., 2008) and FastX toolkit implemented on Galaxy (Bolger et al., 2014). If the read resulting from quality filtering was less than 25 nt in length it was removed. Read mapping and subsequent expression analysis was conducted with the Seqman NGen, and ArrayStar modules of the DNAStar version 16 package (Lasergene, Inc., Madison, WI). Briefly, the *C. elegans* genome assembly WBCel235 (accession number, GCA_000002985.31915) was used as the reference genome for read mapping with the SeqMan NGen transcriptome assembly module. Read mapping was performed with the specifications listed below:

1. mer size (the minimum length of a subsequence - overlapping region of a read fragment required to be considered a match when assembling reads into contigs) = 17.
2. minimum match percentage (specifies the minimum percentage of matches in an overlap that are required to assemble two sequences in a contig) = 95%.
3. minimum aligned length (the minimum length of at least one aligned segment of a read after trimming) = 35.
4. Duplicate sequences were marked and combined.

Gene expression quantitation and statistical analysis of observed differences in gene expression was done with the ArrayStar module. Briefly, normalization of raw read counts was done using the RPKM method (5), and fold change in mRNA abundance between the WT and mutant nDF50 was calculated from total linear RPKM. Significance of observed differences (greater than 2-fold difference in expression,) was determined with the Student’s t-test and the Benjamini-Hochberg correction was employed for multiple hypothesis testing. Genes with significantly altered expression pattern (> 2-fold change; 99% CI), were filtered to retain only those known to express in the *C. elegans* germline. Hierarchical clustering of genes with significantly altered expression in the *C. elegans* germline was performed with the Euclidean distance metric and Centroid linkage method.

### Quantification and statistical analysis

Statistics were run using Prism 7. Statistical details of experiments can be found in the Figure legends. Significance was defined as P < 0.05 for each statistical analysis used.

## DATA AVAILABILITY

The data that support the findings of this study are presented and RNA Sequencing genomic analysis are available at GenBank number XXX.

## RESULTS

### Drosha regulates multiple processes during germline development

To investigate Drosha’s function during germline development we performed two distinct and independent experiments to assess Drosha’s function during germ cell development. a) We analyzed the deletion alleles at different stages of oogenic development and b) we performed a germline-specific RNA interference to deplete Drosha’s activity in adult germlines and assayed dissected germlines for microRNA production as well as germ cell phenotypes. The two existing deletion alleles used were *tm654* and *ok369* (**Figure 1A**) (Materials and Methods). The *C. elegans* Drosha (*drsh-1)* gene contains two Ribonuclease III family domains that process the pri-microRNA into pre-microRNA, as well as the double stranded RNA (dsRNA) binding domain that helps bind with target pri-microRNA (**Figure 1A**). These three domains span exon 6-7 of the eight exons in the gene (**Figure 1A**). The *ok369* deletion removes exon 6 and portions of exon 5 and 7 while the *tm654* deletion removes all of exons 5-8. In both cases, the RNAse IIIb domains are deleted. RT-PCR analysis of the mutants revealed that *ok369* results in an in-frame deletion of the RNAse IIIb domain and part of the dsRNA Binding Domain (**Figure 1A**) while the *tm654* deletion results in a frame shift mutation at amino acid 669, replacing the remaining amino acids with a TGL peptide followed by a stop codon (**Figure 1A**). We performed the germline-specific RNAi analysis rather than an Auxin-induced degron based experiment because we found that tagging Drosha at either the N or the C terminal end with a GFP or AID affects Drosha’s function (Trimmer K.A. and Arur S., unpublished observations).

**Figure 1:**
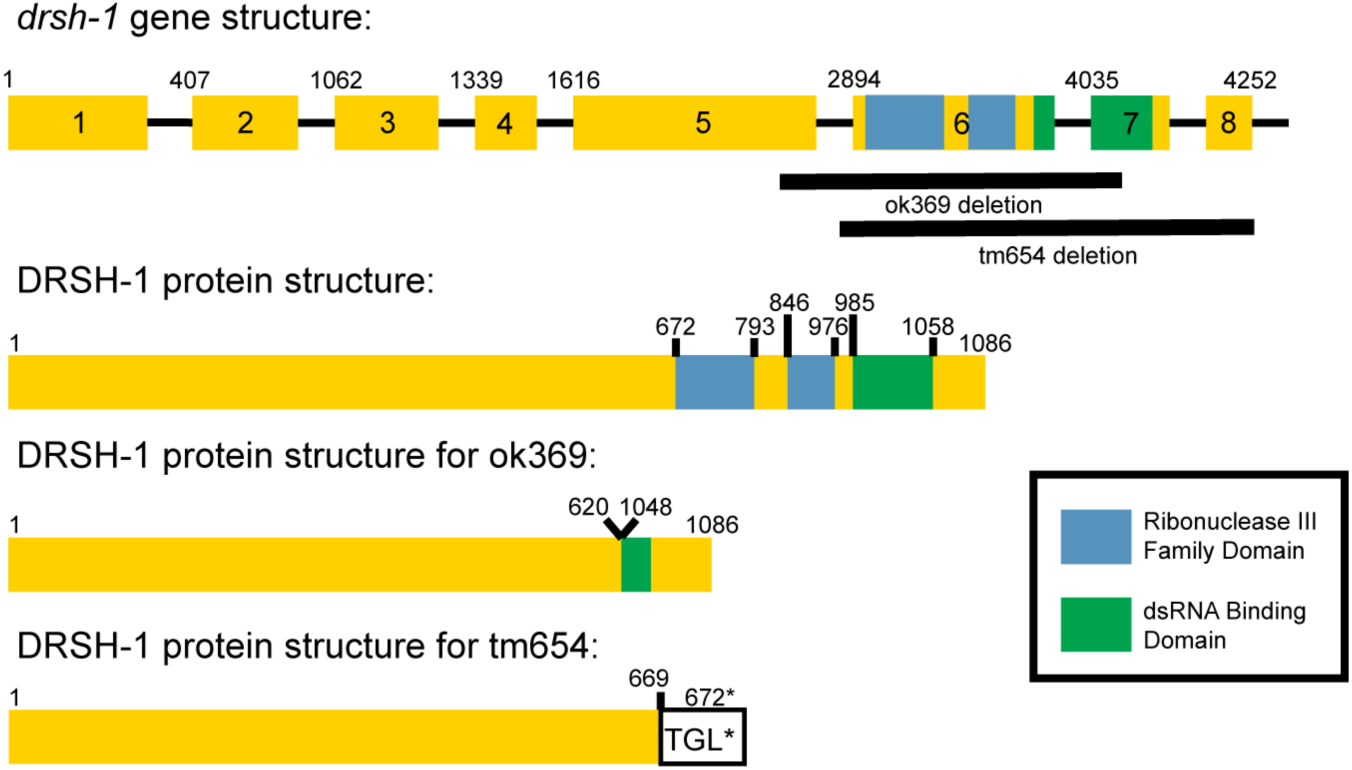
*drsh-1* mutant alleles. Stick diagram of the *drsh-1* gene, deletion breakpoints, catalytic residue (E946), and corresponding protein structure for each allele.

Analysis of the deletion alleles with loss of the RNAse IIIb binding domains in Drosha reveals adult germlines that display endomitotic oocytes, an indicator of failure in meiotic maturation, as described earlier (Minogue et al., 2018; Rios et al., 2017). Meiotic maturation of the terminal oocyte occurs at close to 20 hours past the early L4 stage of the animals, when the hermaphroditic germline has accumulated 3-4 oocytes. Thus, animals that are grown for 18 hours after the early L4 stage carry germlines with oocytes, but do not yet display endomitotic oocytes since meiotic maturation has not been triggered. We utilized this stage of germline development and analyzed dissected germlines from wild type and each of the Drosha deletion alleles for progenitor zone, meiotic region and oocyte formation. We also performed live imaging with transgenic membrane markers (PH domain tagged with GFP and) chromatin markers (mCherry tagged with Histone 2B) in wild type and the deletion alleles to assay for meiotic maturation. In addition, we assayed for germ cell apoptosis using the transgenic marker CED-1 tagged with GFP, which engulfs dying germ cell corpses and can be easily scored in live images.

The distal region of the germline harbors the progenitor zone comprised of both germline stem cells as well as their progenitors (**Figure 2A**). Loss of ALG-1, a homolog of Argonaute 2, has been shown to regulate proliferation in the germline (Bukhari et al., 2012). Thus, we assayed for proliferation using EdU to mark replicating S phase cells and pH3 to mark the dividing M phase cells. We found that at 12 hours past the mid-L4 stage of development (comparable time point to 18 hours past early L4) neither the S phase nor the M phase index were appreciably different in the Drosha deletion alleles compared to wild type animals (**Figure 2B-F**), with S phase index ranging between 65-70% (**Figure 2E**) in the two deletion mutants as well as wild type germlines; M phase index at ∼4 (**Figure 2F**). Although, the total number of germline progenitor cells were significantly reduced in the mutants compared to wild type, suggesting that Drosha regulates the expansion of the progenitor pool (**Figure 2G**). Wild type progenitor zones revealed an average of ∼230 germ cells while the mutants harbored ∼170. This could be due to defects in early larval stage proliferation. Together, these data suggest that Drosha does not regulate proliferation *per se* in the oogenic germline. To determine if Drosha affects the distance from the distal tip cells to where the proliferative cells differentiate, we stained the germlines with HIM-3, which marks the axis of the meiotic chromosomes (Couteau et al., 2004), and is a clear marker for differentiation. Loss of Drosha function results in a reduction in the progenitor zone cell rows to 18 relative to wild type, which enter differentiation at cell rows 20 (**Figure 2H**), suggesting that Drosha may affect the balance of proliferation and differentiation during oogenesis.

**Figure 2:**
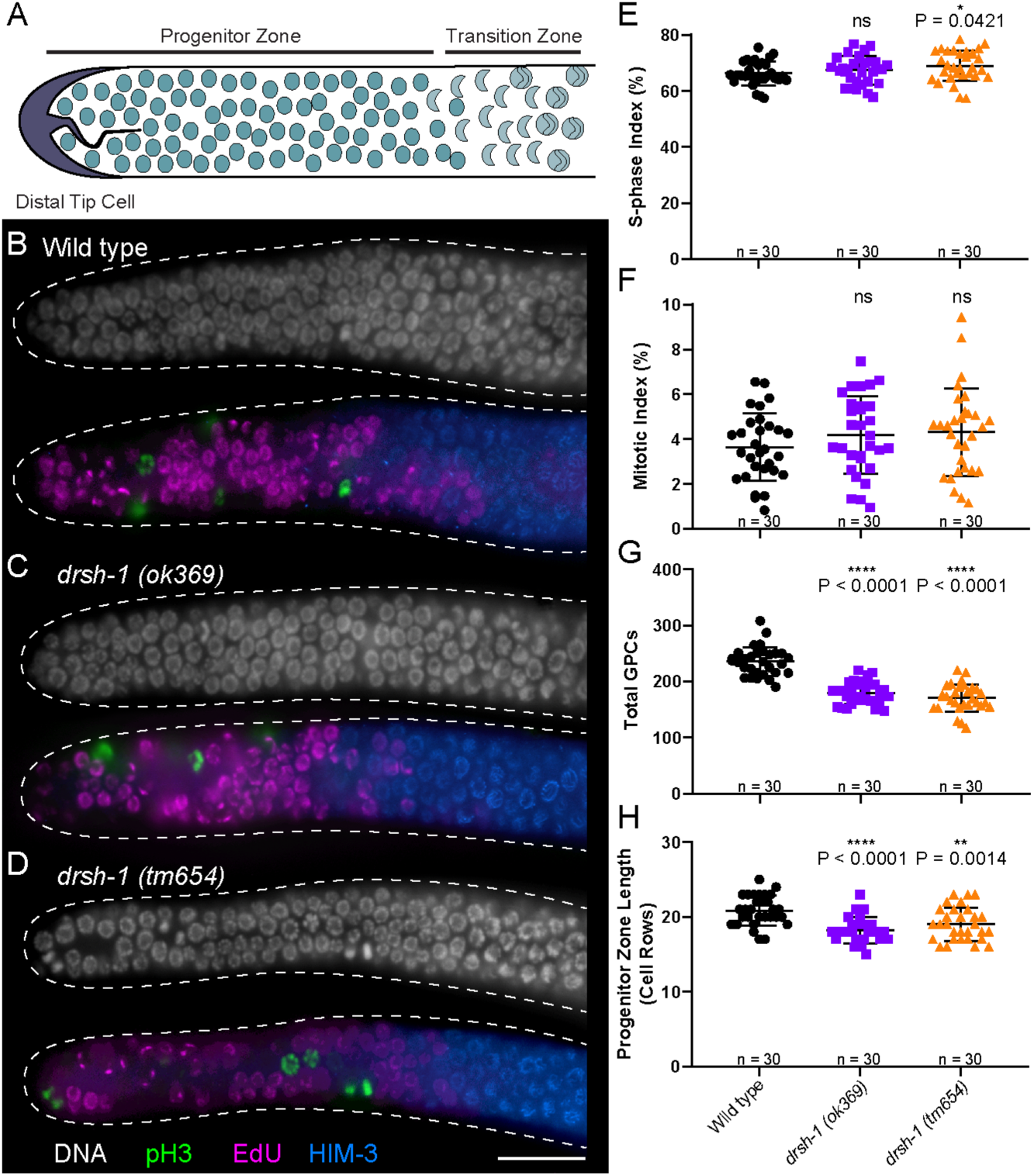
Drosha regulates balance of proliferation and differentiation during oogenesis. **(A)** Schematic view of the adult germline progenitor zone. **(B)** Dissected germline from a wild type animal at 12 hours past mid L4 germlines with **(C)**, Dissected germline from a *drsh-1(ok369)* mutant at 12 hours past mid L4. **(D)** Dissected germline from a *drsh-1(tm654)* mutant at 12 hours past mid L4. **(B**,**C**,**D)** Germlines are labeled for DNA (DAPI, white) pH3 (green), EdU (magenta) and HIM-3 (blue) and are outlined in dashed white. Scale bar = 20µm. **(E)** S phase indices for the indicated genotypes. **(F)** Mitotic indices for the indicated genotypes. **(G)** Total GPCs for the indicated genotypes. **(H)**, Progenitor zone length in cell rows for the indicated genotypes. **(E**,**F**,**G**,**H)** Genotypes, displayed from left to right, are wild type (Black), *drsh-1(ok369)* (Purple), and *drsh-1(tm654)* (Orange). Error bars show 1SD from the mean. (*) p > 0.05, (**) p < 0.01, (****) p < 0.0001 - Statistics were performed using a two-tailed Student’s T-test.

As the germ cells enter meiosis they progress through various stages of meiosis I. To determine whether Drosha regulates different stages of meiosis I progression, we utilized DAPI staining and chromosome morphology to assess whether there were any defects in progression through pachytene. To determine if meiotic progression was affected, we counted the number of pachytene cell rows along the distal to proximal axis of the germline. In wild type, there is an average of 28 pachytene cell rows after which the chromosomes start to remodel at the “loop” region (an anatomical bend of the germline region) and condense their nuclei to form diplotene and diakinetic oocytes (**Figure 3A-B**). In both Drosha deletion alleles however, the average number of pachytene cell rows is significantly higher with an average of 32 cell rows that continue past the loop region in each mutant (**Figure 3A-B**). These data suggest that the pachytene cells do not efficiently transition into diplotene oocytes at the loop region, but instead linger in pachytene past the loop region of the germline (**Figure 3A-B**). The outcome of germ cell differentiation is the production of individualized oocytes. The lack of effective transition of pachytene stage nuclei into diplotene and diakinesis resulted in an overall reduction in the number of individualized oocytes in the *drsh-1* mutants (**Figure 3C**). Collectively, these data suggest that Drosha functions to promote pachytene progression and oocyte development. We had previously identified two Drosha-dependent microRNAs that accumulate in late pachytene and developing oocytes, miR-35 and miR-61. Loss of either of these microRNAs *via* deletion mutants or RNAi of the pre-microRNAs results in pachytene progression defects (Minogue et al., 2018), leading to the model that Drosha may regulate pachytene progression and oocyte development through miR-35 and miR-61.

**Figure 3:**
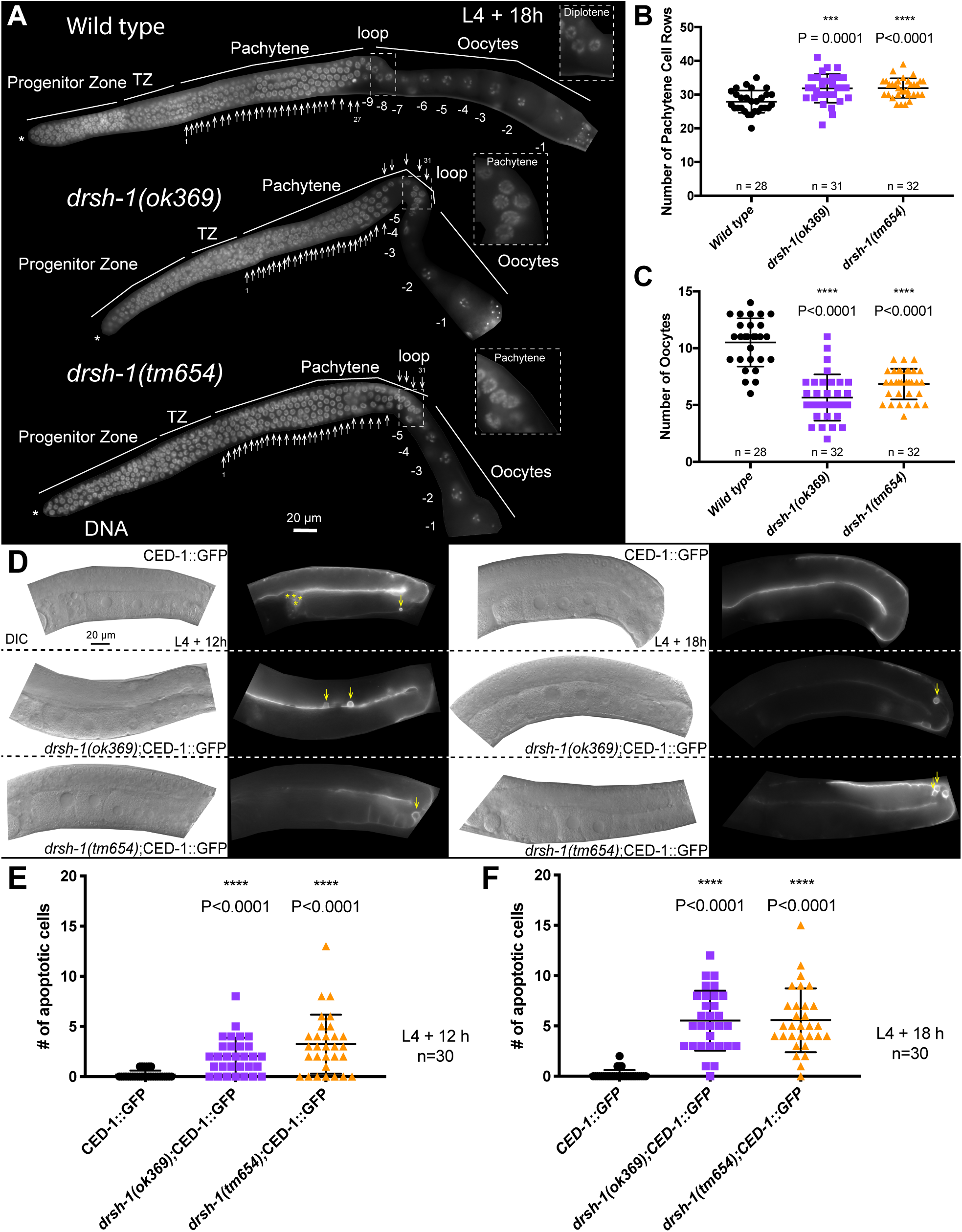
Drosha regulates meiotic processes. **(A)** Dissected germlines from young adult L4 +18 h hermaphrodites labeled with DAPI (white, DNA). White (*) Denotes the distal end of each germline. Arrows point to each counted pachytene row. Inset displays chromosomal morphology at the loop region of the germline. Oocytes are numbered starting from the oocyte closest to the spermatheca, termed as the −1 oocyte and counted onwards. **(B)** Graph displaying the number of pachytene cell rows from described genotypes at L4+18 h. **(C)** Graph displaying the total number of oocytes from described genotypes at L4+18 h. **(D)** Live imaging of CED-1::GFP, *drsh-1(ok369)*;CED-1::GFP, and *drsh-1(tm654)*;CED-1::GFP hermaphroditic animals at L4+12 h and L4+18 h. Images were acquired as Z-stacks with a representative slice shown here for each genotype. Arrows point to apoptotic cells marked with CED-1::GFP. Yellow (*) indicates sperm residual bodies, which were not counted. **(E)** Graph displaying the number of apoptotic cells from the described genotypes at L4+12 h. **(F)** Graph displaying the number of apoptotic cells from the described genotypes at L4+18 h. In (B-C, E-F) error bars indicate mean ±SD. Statistical significance was calculated by a Mann-Whitney test.

As germ cells progress through pachytene, a significant population undergoes physiological apoptosis (Gartner et al., 2008). To determine whether Drosha regulates germ cell apoptosis, we crossed in the CED-1::GFP (Zhou et al., 2001) transgenic line that marks somatic sheath cells engulfing early apoptotic corpses into the two Drosha deletion alleles. Live imaging with Z-stacks was used to quantify the number of apoptotic cells at both 12 and 18 hours after the L4 stage of development (**Figure 3D**). On an average, wild type germlines display few or no apoptotic germ cells at any one-time point. Loss of Drosha function however, led to a significant increase in the apoptotic germ cells (**Figure 3E-F**). miR-35 family has been previously shown to regulate germ cell apoptosis *via* antagonizing the BH3-domain containing gene *egl-1* (Tran et al., 2019). Together, these data suggesting that Drosha may normally function to inhibit germ cell apoptosis in a miR-35 dependent manner.

Loss of Drosha in adults produces endomitotic oocytes, which is a hallmark of meiotic maturation defects (Minogue et al., 2018; Rios et al., 2017). To determine when during meiotic maturation Drosha mutant oocytes start to present with defects, we performed live imaging of wild type and Drosha meiotic maturation using transgenic lines with PH domain tagged GFP and histone tagged with mCherry. Each oocyte matures and ovulates into the spermatheca, once every ∼ 23 minutes, where it completes meiosis I and II prior to undergoing fertilization (Greenstein, 2005). Upon completion of meiosis and fertilization, the newly formed zygote exits the spermatheca and enters the uterus where embryogenesis begins.

We performed time lapse imaging of wild type and Drosha mutant worms with four minute intervals for 60-90 minutes total. During this analysis, we observed maturation and ovulation of the oocyte into the spermatheca followed by the ovulated and fertilized oocyte exiting into the uterus (**Figure 4A**). In both of the Drosha deletion alleles, much like wild type animals, we observed that the first few oocytes mature into the spermatheca and are fertilized; an example of a fertilized embryo in the uterus is shown (**Figure 4A**). However, subsequent oocytes complete maturation but fail to ovulate (**Figure 4A**). These data suggest that the Drosha mutant oocytes are themselves competent to mature and ovulate even though they are slightly delayed in the process, suggesting that the emergence of the endomitotic oocytes in younger animals may be due to a feedback from the soma where the embryos are arrested and do not progress. The embryos from both Drosha mutant alleles are misshapen and do not progress beyond the 4-cell stage, and eventually the embryos display polyploid oocytes (**arrow, Figure 4B**). Overall, these data suggest that the transition of an oocyte to a totipotent embryo is regulated by Drosha.

**Figure 4:**
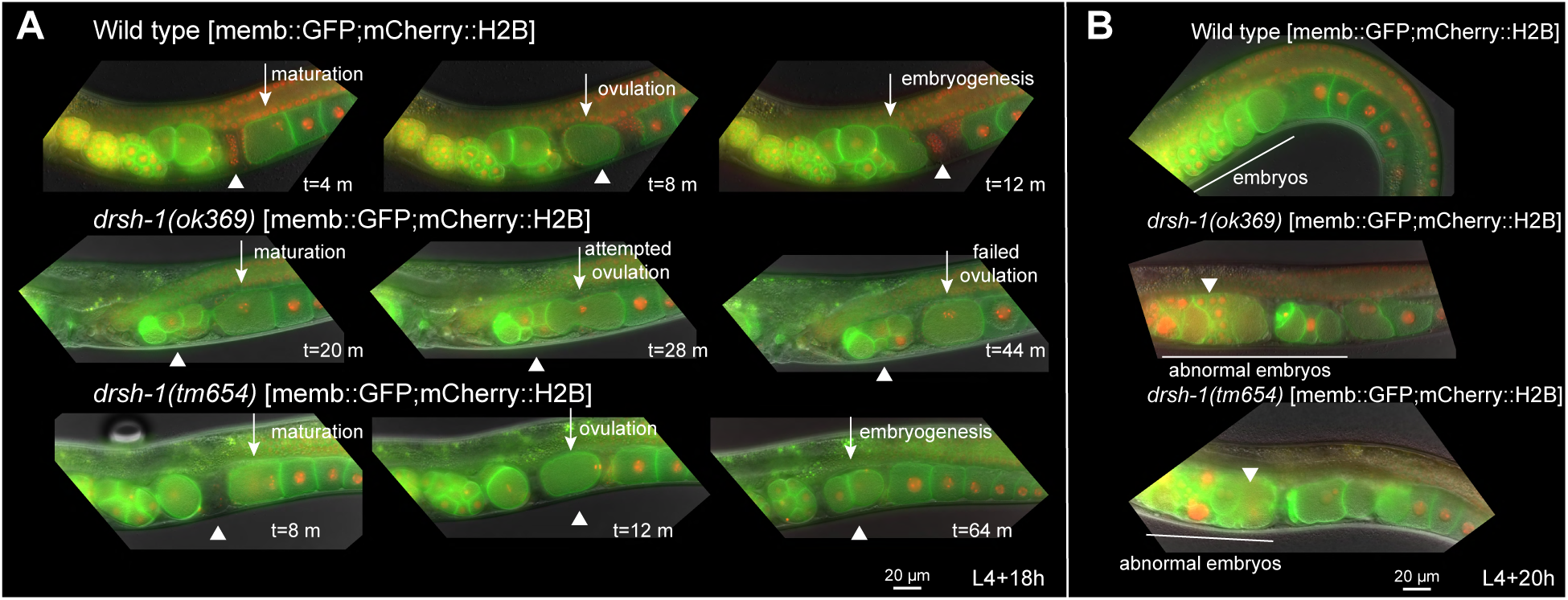
Drosha regulates oocyte-to-embryo transition. **(A)** Live imaging performed on animals of indicated genotypes at L4+18 hours hermaphrodites. Membrane GFP; mCherry::H2B wild type animals show oocytes that mature, ovulate four minutes later, undergo fertilization, and then exit the spermatheca for embryogenesis. In both the *drsh-1* deletion mutants, early oocytes mature and undergo ovulation, and early embryogenesis. **(B)** Live imaging performed on animals of indicated genotypes at L4+20 hours hermaphrodites. At this stage in development, the early oocytes have matured and ovulated and fail to undergo normal embryogenesis. The oocytes in *drsh-1* mutants develop into abnormal embryos that are malformed and misshaped relative to wild type. Scale bar: 20 μm

In a mutant hermaphroditic germline, the embryonic defects can also arise due to defects in sperm development. To determine whether the oocyte-to-embryo defects were sperm dependent, we mated wild type males to Drosha mutant lines harboring transgenic PH domain tagged with GFP and Histone tagged with mCherry (Methods). In these lines, the hermaphroditic sperm is red in color, thus we can distinguish between the unlabeled sperm emanating from the male *vs* the labeled sperm from the hermaphrodite. First, we asked whether mating with wild type males could rescue the sterility of the Drosha deletion mutants. While mating with wild type males led to an increase in progeny in wild type hermaphrodites, (**Figure S1A**), wild type males did not rescue the sterility in the Drosha mutant hermaphrodites (**Figure S1A**). Careful examination of the spermatheca from the mated animals revealed the presence of unlabeled sperm (**Figure S1B-D**) in the Drosha mutants, and the mutants still displayed endomitotic oocytes and abnormal embryos in the presence of the wild type sperm, suggesting that the underlying cause of sterility in Drosha mutants may not be sperm dependent (**Figure S1B-D**). These analyses together demonstrate that Drosha regulates pachytene progression, germ cell apoptosis and oocyte-to-embryo transition during oogenesis. Interestingly, these phenotypes are consistent with the expression of the small repertoire of Drosha-dependent oocyte-expressed microRNAs (Minogue et al., 2018). To determine whether Drosha regulates the oogenic phenotypes and microRNA expression in a germline autonomous manner, we assayed for germline autonomous functions of Drosha.

### Drosha regulates pachytene progression and oocyte development germline autonomously through miR-35 and miR-51

To determine the functions of *drsh-1* that are germline autonomous, we performed a germline-specific RNAi experiment (Method). We utilized germline-specific RNAi depletion since tagging of GFP or AID to Drosha N or C terminal ends led to loss of Drosha function in our hands (Trimmer K.A. and Arur S., unpublished observation). To perform germline-specific depletion of Drosha, we took advantage of a recently published reagent, wherein the worms have a null mutation in the Argonaute *rde-1*, which is necessary for RNAi (Firnhaber and Hammarlund, 2013), combined with a functional germline-expressed RDE-1 transgene (Zou et al., 2019). In this strain, because RDE-1 is expressed only in the germline, RNAi is executed only in the germline and allows for germline specific depletion of the gene of interest.

We performed feeding RNAi (Methods) on the germline-specific RNAi line against Drosha (*drsh-1*). Luciferase RNAi was used as a negative control and *glp-1* RNAi, the loss of which causes a severe reduction in germline proliferation (Crittenden et al., 2003; Kimble and Simpson, 1997; Seydoux and Schedl, 2001). Levels of Drosha depletion was assessed *via* qRT-PCR on the dissected germlines as compared to control RNAi (**Figure 5A**). To obtain significant reduction of RNA to be able to confidently assess the phenotypes, we had to perform an F2 RNAi, which led to ∼75% reduction in Drosha expression as assessed by qRT-PCR analysis (**Figure 5A**).

**Figure 5:**
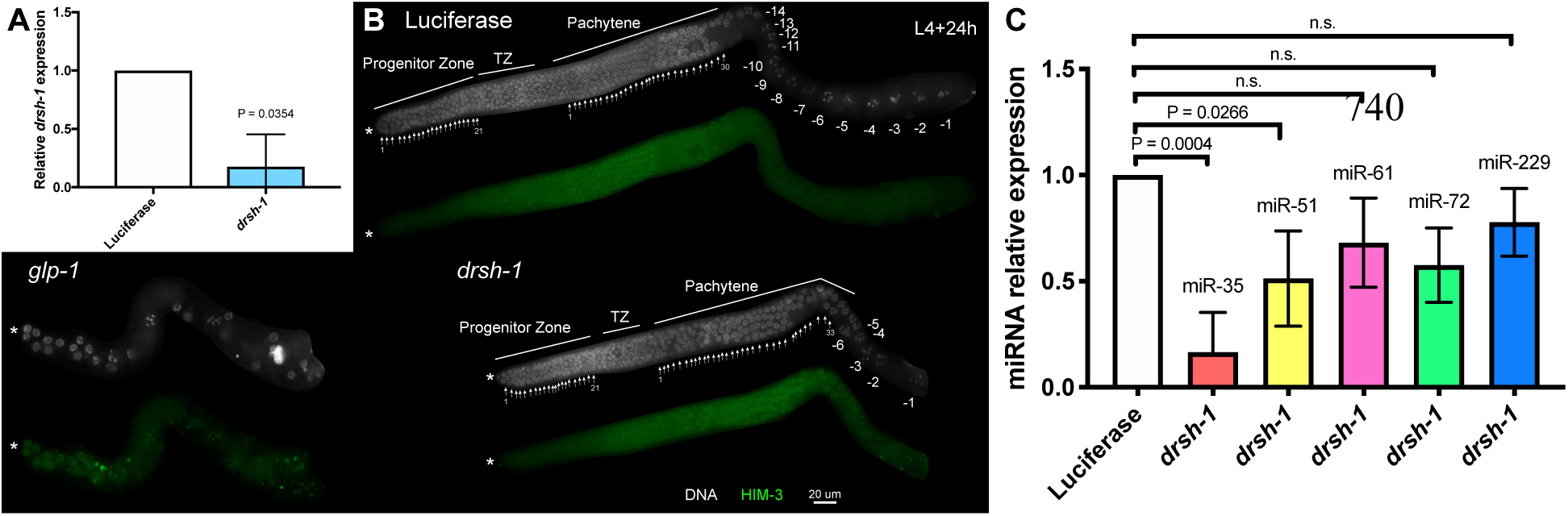
Drosha regulates pachytene progression, oocyte development, miR-35 and miR-51 germline autonomously. **(A)** qRT-PCR analysis for *drsh-1* mRNA on dissected germlines from control (luciferase RNAi) and *drsh-1* RNAi F2 animals averaged across three independent replicates. (**B)** Dissected germlines from adult L4+24 hours hermaphrodites labeled with DAPI (white, DNA) and HIM-3 (green, meiotic germ cells). White (*) denotes the distal end of each germline. Arrows point to each counted proliferative zone row or pachytene row. **(C)** TaqMan analysis of relative microRNA knockdown from each RNAi treatment group. miR-35 and miR-51 are significantly reduced in *drsh-1* depleted germlines compared to the Luciferase RNAi control germlines. TaqMan assay was repeated for three independent replicates, and statistical significance was calculated by a one tailed ANOVA with Bonferroni correction. Mean ±1SD is graphed. Scale bar: 20 μm

We analyzed the F2 RNAi germlines at 24 hours past the L4 stage of development and assessed phenotypes identified in the deletion mutants using DAPI and HIM-3. Unlike in the deletion mutants which displayed lower germline progenitor cell numbers and progenitor zone length (**Figure 2**), we found that these aspects of the progenitor zone of Drosha depleted germlines were similar to wild type (**Figure 5**). We next assessed for oogenic phenotypes.

Similar to what we observed in the Drosha deletion mutants, we found that germline specific depletion of Drosha led to pachytene progression defects and reduced number of oocytes (**Figure 5B**), while control Luciferase RNAi germlines displayed wild type germline development (**Figure 5B**) and *glp-1* RNAi presented with a strong glp phenotype (**Figure 5B**). However, the Drosha depleted oocytes while lower in number, however did not enter endomitosis (**Figure 5B**), suggesting that as observed in the Drosha deletion mutants, meiotic maturation was not regulated by Drosha in a germline autonomous manner. These data demonstrate that Drosha regulates pachytene progression and oocyte development germline autonomously, and oocyte meiotic maturation in a germline non-autonomous manner. We next assayed for Drosha-dependent oocyte-expressed microRNAs to determine which ones were acting germline autonomously to regulate these phenotypes.

We assayed two Drosha-dependent microRNAs that regulates pachytene progression and two that regulate oocyte development (**Table 1**). miR-35 and miR-61 accumulate in late pachytene and into the developing and arrested oocytes in a Drosha-dependent manner (Minogue et al., 2018). To determine whether miR-35 and miR-61 both function germline autonomously to regulate pachytene progression, we assayed for their expression in the Drosha depleted germlines using TaqMan based analysis. miR-35 was significantly reduced upon Drosha depletion in the germline (**Figure 5C, red bar**), however miR-61 was not (**Figure 5C, pink bar**). These data were surprising, and suggested the pachytene progression defect in Drosha mutants could be phenocopied by loss of miR-35 alone in a germline autonomous manner. miR-61 may accumulate in a Drosha-dependent manner when generated in the soma, but is not germline autonomous in its generation.

**Table 1:** Germline expression, and autonomony of Drosha-dependent oocyte-expressed microRNAs.

| microRNA | Germline Expression | Germline Phenotype | Germline Autonomy |
| --- | --- | --- | --- |
| *mir-35* | Pachytene, Oocytes | Pachytene Progression | Germline Autonomous |
| *mir-61* | Pachytene, Oocytes | Pachytene Progression | Germline Non autonomous |
| *mir-51* | Oocytes | Fewer Oocytes | Germline Autonomous |
| *mir-72* | Oocytes | Fewer Oocytes | Germline Non autonomous |

miR-72 and miR-51 accumulate in diakinetic oocytes in a Drosha-dependent manner (Table 1). While reduction of either microRNA is sufficient to result in fewer oocytes relative to wild type, miR-72 is not significantly reduced in the germline upon Drosha depletion but miR-51 is (**Figure 5C**). Oocyte development is thus controlled by miR-51 and miR-72 in both a germline autonomous and non-autonomous manner. These data suggest that Drosha regulates events of pachytene progression and oocyte development in a germline autonomous manner, while progenitor zone dynamics and meiotic maturation are regulated in a germline non-autonomous manner. We also find that though Drosha-dependent oocyte-expressed microRNAs, miR-61 and miR-72, are sufficient to regulate pachytene progression and oocyte development respectively, they are generated in a germline non-autonomous manner.

### The miR-35 family regulates germline functions likely through translational inhibition of targets

Given that the miR-35 family lies downstream to *drsh-1* and functions germline autonomously for pachytene progression and oocyte development, we next wanted to determine what potential genes this family of microRNAs may be regulating. To determine any potential mRNA targets, we extracted RNA from whole animals from wild type and *miR-35-41(nDf50)* mutants and performed RNA sequencing analysis in three independent replicates (Methods). Because this analysis was performed on whole animals, we first identified a set of 469 differentially expressed genes in the *miR-35-41(nDf50)* mutants compared to wild type (**Table S1, Figure 6**). To narrow the list down to potential targets during oogenesis, we cross-referenced our list to a published set of germline enriched genes in *C. elegans* (Reinke et al., 2004). This analysis led us to 41 differentially expressed genes identified to be either germline intrinsic or oocyte specific (**Table S2**). To narrow the list down further, we searched for known expression pattern data from NEXTDB Ver.4.0 (Kohara, 2001) and WormBase (WS273) (**Table S2**). However, the majority of genes we identified still have unknown expression patterns within the germline. Moving forward with the full list of 41 genes, we surmised that any genes upregulated in the mutants compared to wild type could be potential mRNAs degraded by the miR-35 family, and any genes downregulated in the mutants compared to wild type could be potential mRNAs protected by the miR-35 family. Of the 41 differentially expressed genes, 39 genes were significantly upregulated (**Table S2**), while only two were significantly downregulated (**Table S2**). 33 genes were higher in the mutant than wild type, but at 1.5 fold. To determine if the miR-35 family can bind the 3’ UTR of the 41 potential targets, we used TargetScanWorm 6.2 (Jan et al., 2011; Lewis et al., 2005) to assess presence of seed sequences. Interestingly, none of the 41 mRNAs contain the miR-35 seed match within their 3’ UTR (**Table S2**), suggesting that these genes may not be regulated by miR-35 through canonical seed sequence matches. It is possible that more detailed characterization of 3’ UTRs may help identify seed sequences in mRNA targets. However, the current result is consistent with recent studies that suggest that microRNAs may regulate targets in a manner distinct from the canonical seed sequence (Chipman and Pasquinelli, 2019; Nicholson and Pasquinelli, 2019).

**Figure 6:**
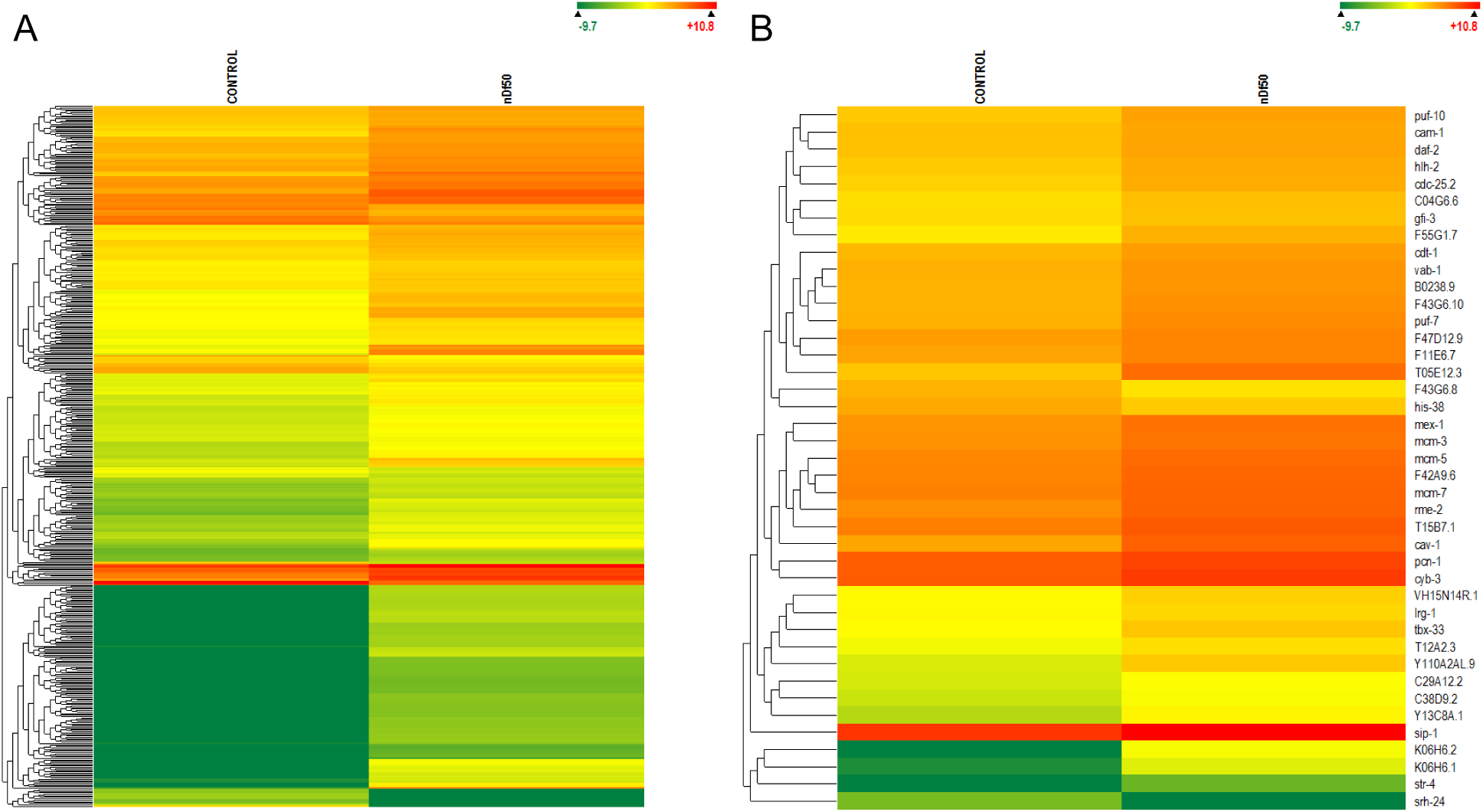
RNA Sequencing analysis of wild type and *mir-35-41* mutant animals. RNA Sequencing analysis from wild type and nDf50 whole animals reveals only 496 RNAs that are significantly changed (CI 99%, **A**), of which only 41 are germline genes (**B**).

**Figure 7:**
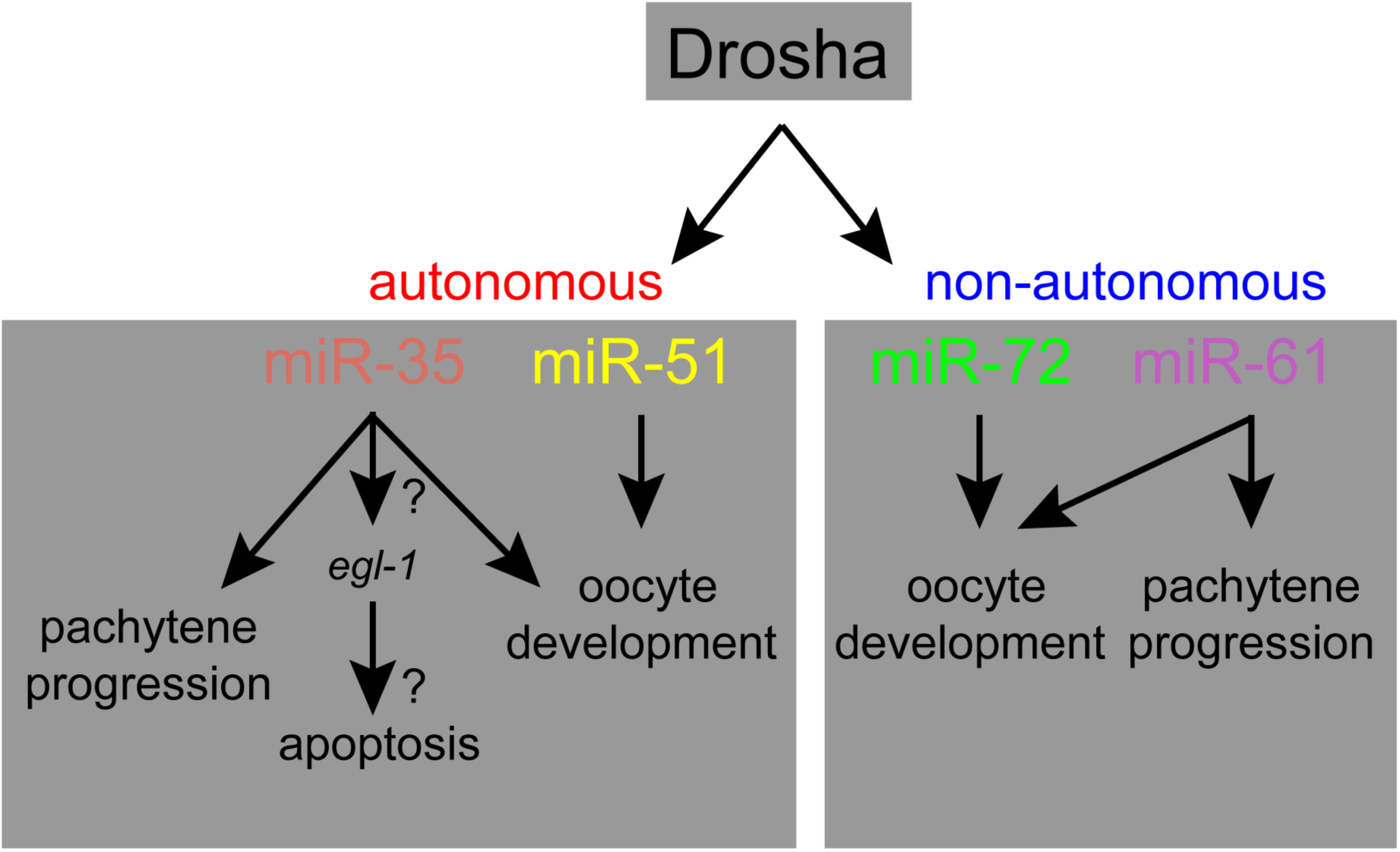
Model. Drosha-dependent oocyte-expressed microRNA circuit coordinates oocyte development in a germline autonomous manner as well as through soma germline communication.

We scanned the literature to assess whether gain of any of the 39 genes was implicated in pachytene progression or oocyte development defects. Interestingly, for some genes such as *daf-2* and *mex-5* loss, rather than gain, has been shown to regulate germ cell development (Michaelson et al., 2010; Schubert et al., 2000). That *daf-*2 and *mex-5* may be repressed during normal development by miR-35 is counterintuitive, raising the possibility that miR-35 functions to balance the level of the gene rather than completely turn it off. In this manner miR-35 would be able to titrate gene expression rather than regulate it in a switch-like manner may help lend physiological robustness to the germline. It is also likely that the pachytene progression and oocyte development defect may be regulated by miR-35 in a translational manner rather than through post transcriptional degradation. Overall, the miR-35 family regulates germline mRNAs through a non-canonical manner and may function to maintain the robustness and fidelity of the germline.

## DISCUSSION

A role for microRNAs and microRNA pathway genes in regulating oocyte development has been widely debated in recent years, in part because of divergent results between mammalian models and invertebrate models. For example, in mice, oocyte specific loss of Dgcr8 or Pasha in mammals does not yield oocyte defects (Suh et al., 2010); loss of Dicer leads to defects in oocyte maturation in mammals, and this function is attributed to endo-siRNA function (Stein et al., 2015) rather than microRNAs. In worms, on the other hand, oocyte meiotic maturation was attributed to loss of Dicer, Drosha and Pasha, since loss of each causes endomitosis of oocytes (Knight and Bass, 2001; Minogue et al., 2018; Rios et al., 2017). However, we showed that Dicer functions germline non-autonomously to regulate meiotic maturation through regulation of endo-siRNAs (Drake et al., 2014), a result that was found to be true in mammalian biology as well (Stein et al., 2015). We then assayed for oocyte-expressed microRNAs, and found that a very small repertoire of microRNAs is oocyte-expressed, and about only half of these are Drosha-dependent (Minogue et al., 2018). In this study, we delineate the function of Drosha during germline and oocyte development by using two deletion alleles and germline-specific depletion. We show that Drosha regulates the transition of pachytene cells to diplotene during meiosis I in a germline autonomous manner, as well as through non-autonomous manner. In this role, miR-35 functions germline autonomously, while miR-61 functions germline non-autonomously. In addition, Drosha regulates oocyte-to-embryo transition in a germline non-autonomous manner. We find that Drosha does not present with a defect in oocyte meiotic maturation *per se*, and the endomitotic phenotype may be attributed to a potential feedback regulation from the uterus and soma into the germline. This study helps resolve the controversy between mammalian and invertebrate fields on the role of microRNA pathway genes during meiotic maturation, while highlighting a role for the microRNA pathway and the miR-35 family and miR-61 in progression of meiosis I. It is very likely that a small proportion of microRNA families similarly regulates meiotic progression in mammals.

The Drosha-dependent oocyte-expressed microRNAs have been mapped *via in situ* hybridization technology to different expression patterns (Minogue et al., 2018), for example, Pattern 1 microRNAs accumulate from late pachytene and into the developing or arrested oocytes, while Pattern 2 or 3 microRNAs accumulated in developing or arrested oocytes. While these microRNAs are expressed in distinct patterns and accumulate in one oocyte but not its neighboring oocyte, the question remained whether the microRNAs were generated in the oocyte, or whether they traversed the soma to accumulate in the oocyte. This is a pertinent question since small RNAs can traverse tissues and execute their function in cell types distinct from the cells that generate the RNAs. In addition, because oocytes are transcriptionally silent, a formal possibility remained that the microRNAs in the oocytes were obtained from the soma, while those generated in late pachytene could be germline autonomous. These possibilities were complemented by recent studies that have debated whether small RNAs under the size of 50 bp can be taken up by the yolk receptor RME-2 (Antony Jose, personal communication) or the small RNA transporter SID-1 (Wang and Hunter, 2017). We found that indeed some of the microRNAs that are oocyte-expressed are germline non-autonomous, since depletion of Drosha specifically from the germline did not affect their accumulation in the dissected germlines. Interestingly, however, these belong to both Pattern 1 and 2 microRNAs. For example, miR-61 expressed from late pachytene into the developing oocytes (Minogue et al., 2018) and miR-72 which belongs to Pattern 2 (Minogue et al., 2018) and accumulates only in arrested oocytes, are both generated by Drosha in a germline non-autonomous manner. This is interesting because miR-35 mimics miR-61 in expression and miR-51 mimics miR-72 in expression and both miR-35 and miR-51 are germline autonomous. Thus, why some microRNAs are produced in a cell or tissue autonomous manner while others are not cannot be commented on simply based on their expression pattern. This study starts to delineate the function and cell of action for microRNAs during oogenesis.

Consistent with a role for Drosha during pachytene progression, miR-35 loss also regulates pachytene progression and chromosome transition during oogenesis. The miR-35 family has been shown to regulate only translational inhibition in the germline *vs* post-transcriptional degradation (Dallaire et al., 2018). Deep sequencing the miR-35 *nDf50* mutant reveals a small subset of germline mRNAs that are significantly upregulated in the mutant relative to wild type controls. However, none of these mRNAs bear the seed sequence for miR-35. These observations suggest three possibilities (a) that these targets are upregulated several fold suggests that miR-35 may be able to regulate post transcriptional degradation, but may do so in a novel manner that may not be seed sequence dependent as has been suggested recently in literature for microRNAs (Chipman and Pasquinelli, 2019; Nicholson and Pasquinelli, 2019), (b) that longer 3’UTR’s may carry the miR-35 seed sequence, or (c) it is likely that miR-35 translationally inhibits a particular target gene, which may in turn cause transcriptional de-repression in the germline, and in this manner, miR-35 may regulate the mRNAs indirectly. Future studies should be able to delineate these possibilities. None of the germline genes that were upregulated in the miR-35 mutant seemed potentially to be candidates that could regulate pachytene progression, in part because these genes are normally not de-repressed in the germline, for example like *daf-2* and *mex-5*. However, it is likely that these are spatially titrated across the germline either post-transcriptionally or translationally by the miR-35 cluster to maintain a stoichiometric balance and lead to robustness in germ cell development. Thus, Drosha-dependent microRNAs seem to regulate events of oogenesis in a germline autonomous manner as well as through germline and soma communication and maintain germline robustness and fidelity.

## ACKNOWLEDGMENTS

We thank members of the Arur Lab for critical discussions throughout this study. MD Anderson DNA Sequencing Facility conducted the RNA Sequencing of the miR-35 mutants supported by the NCI Cancer Center Support Grant.

