## Supplemental Figures and Legends for "Drosha Regulates Oogenesis and microRNAs Germline Autonomously and Non-autonomously in *C. elegans*"

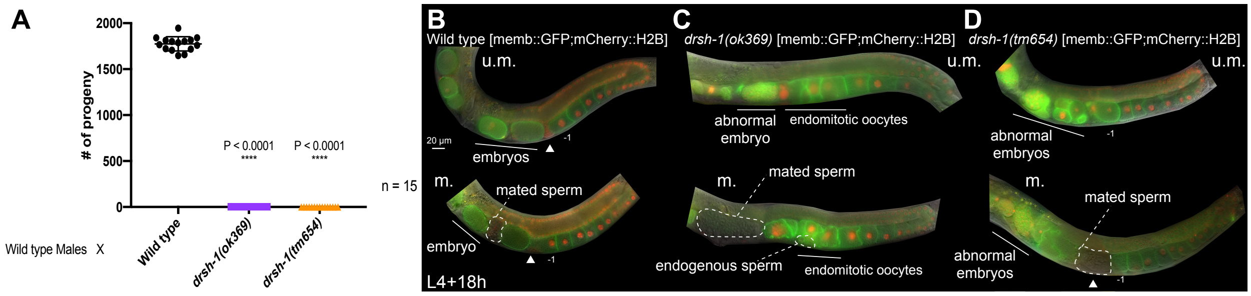


**Figure S1: Wild type sperm cannot rescue the *drsh-1* fertility and oocyte-to-embryo defects. (A)** *drsh-1* mutants produce no progeny even when mated to wild type males. **(B)** Live imaging acquired at L4+18 h from memb::GFP; mCherry::H2B hermaphrodites that are either unmated (u.m.) or mated (m.) to wild type males. Hermaphrodite sperm is labeled with mCherry::H2B while wild type mated sperm is unlabeled. In either the unmated or mated case, memb::GFP; mCherry::H2B hermaphrodites produce embryos. **(C)** Live imaging acquired at L4+18 h from *drsh-1(ok369);* memb::GFP; mCherry::H2B hermaphrodites that are either unmated (u.m.) or mated (m.) to wild type males. In either the unmated or mated case, *drsh-1(ok369);* memb::GFP; mCherry::H2B mutants are still unable to produce wild type embryos and display an oocyte-to-embryo defect. **(D)** Live imaging acquired at L4+18 h from *drsh-1(tm654);* memb::GFP; mCherry::H2B hermaphrodites that are either unmated (u.m.) or mated (m.) to wild type males. In either the unmated or mated case, *drsh-1(tm654);* memb::GFP; mCherry::H2B mutants are still unable to produce wild type embryos and display an oocyte-to-embryo defect.

**SUPPLEMENTAL TABLE:**

**Table S1: Genes significantly up or down regulated in *mir-35-41(nDf50)* mutants.** All genes are listed based on a 99% confidence interval.

**Table S2: Genes significantly up or down regulated in miR-35-41(nDf40) mutant sorted by germline enrichment and expression.**
